# BioChemAIgent: An AI-driven Protein Modeling and Docking Framework for Structure-Based Drug Discovery

**DOI:** 10.64898/2025.12.17.694892

**Authors:** Behnam Yousefi, Nora Constanze Laubach, Sven Heins, Lucia Testa, Søren W. Gersting, Stefan Bonn

**Affiliations:** Institute of Medical Systems Bioinformatics, Center for Biomedical AI (bAIome), Center for Molecular Neurobiology (ZMNH), University Medical Center Hamburg-Eppendorf, Hamburg, 20251, Germany; Hamburg Center for Translational Immunology (HCTI), University Medical Center Hamburg-Eppendorf, Hamburg, 20251, Germany; German Center for Child and Adolescent Health (DZKJ), partner site Hamburg, University Medical Center Hamburg-Eppendorf, Germany; University Children’s Research, UCR@Kinder-UKE, University Medical Center, Hamburg-Eppendorf, Hamburg 20251, Germany

## Abstract

Recent advances in AI have substantially accelerated in silico drug discovery, yet most computational approaches remain task-specific and require expert knowledge. We present **BioChemAIgent**, an agentic framework that orchestrates state-of-the-art AI models and established computational chemistry tools to support end-to-end small-molecule analysis, protein modeling, molecular docking, and interaction analysis through a unified interface. BioChemAIgent emphasizes transparent reasoning, reproducible workflows, and community-oriented extensibility for structural biology and drug discovery applications.

## Main

Traditional drug discovery is a lengthy, expensive, and failure-prone process that can take over a decade and billions of dollars to bring a single therapy to market ^1^. AI-based methods have begun to reshape this landscape, accelerating key stages from protein structure prediction to binding pose and affinity estimation of drug candidate compounds ^2–5^. Yet, despite rapid methodological progress, most computational tools remain optimized for narrowly defined tasks, resulting in a fragmented ecosystem that demands expert integration for practical use. Molecular docking, for instance, requires the sequential integration of multiple specialized tools for both protein and ligand preprocessing, e.g. protonation and energy minimization, as well as post-processing for protein–ligand interaction analysis and visualization.

Recent advances in large language models (LLMs) and agent-based systems, such as model context protocol (MCP) ^6^ and agent2agent (A2A) protocol, are reshaping what is technically possible in computational science. These systems allow models to directly interact with tools and datasets and reason over available information, which enables them to plan multi-step workflows and dynamically select and run the most suitable tools for a given task. At the same time, the agents offer a simple and intuitive user experience, enabling broad access regardless of technical expertise. General-purpose agents still struggle in tasks that need deep, domain-specific expertise, and cannot reason across specialized domains like structural biology or cheminformatics ^7,8^. Hence, complex domains, such as drug discovery, require an AI agent that is aware of existing computational methods, their strengths, limitations, and proper use, and can assemble them into coherent, transparent and reproducible workflows–improving both scientific rigor and output quality.

Here, we present BioChemAIgent, an agentic system that orchestrates state-of-the-art AI models and established computational chemistry tools for end-to-end drug discovery applications. The platform provides a unified interface for key drug development tasks like small molecule analysis, protein modeling and molecular docking, including protein-ligand interaction analysis, and molecular visualization (Fig. 1). By integrating best-practice workflows, BioChemAIgent bridges the gap between powerful predictive models and a readily executable, reproducible drug discovery workflow. It has been equipped with curated documentations that support dynamic communication between tools and models. This design is inspired by the coordinated interplay of cross-functional expert teams, a core principle that enables BioChemAIgent to reproduce interdisciplinary workflows and empower lean teams with capabilities typically requiring broader expert groups. Besides, this study provides a community-oriented registry specialized for the development of agentic systems for drug discovery and structural biology, openly accessible on Github ^i^. Domain experts can bring forward ideas to be implemented, and developers can propose or refine MCP servers to broaden and improve the agent’s functionality. We further provide a publicly accessible web interface ^ii^with both a chatbot and a molecular structure viewer.

**Figure 1.**
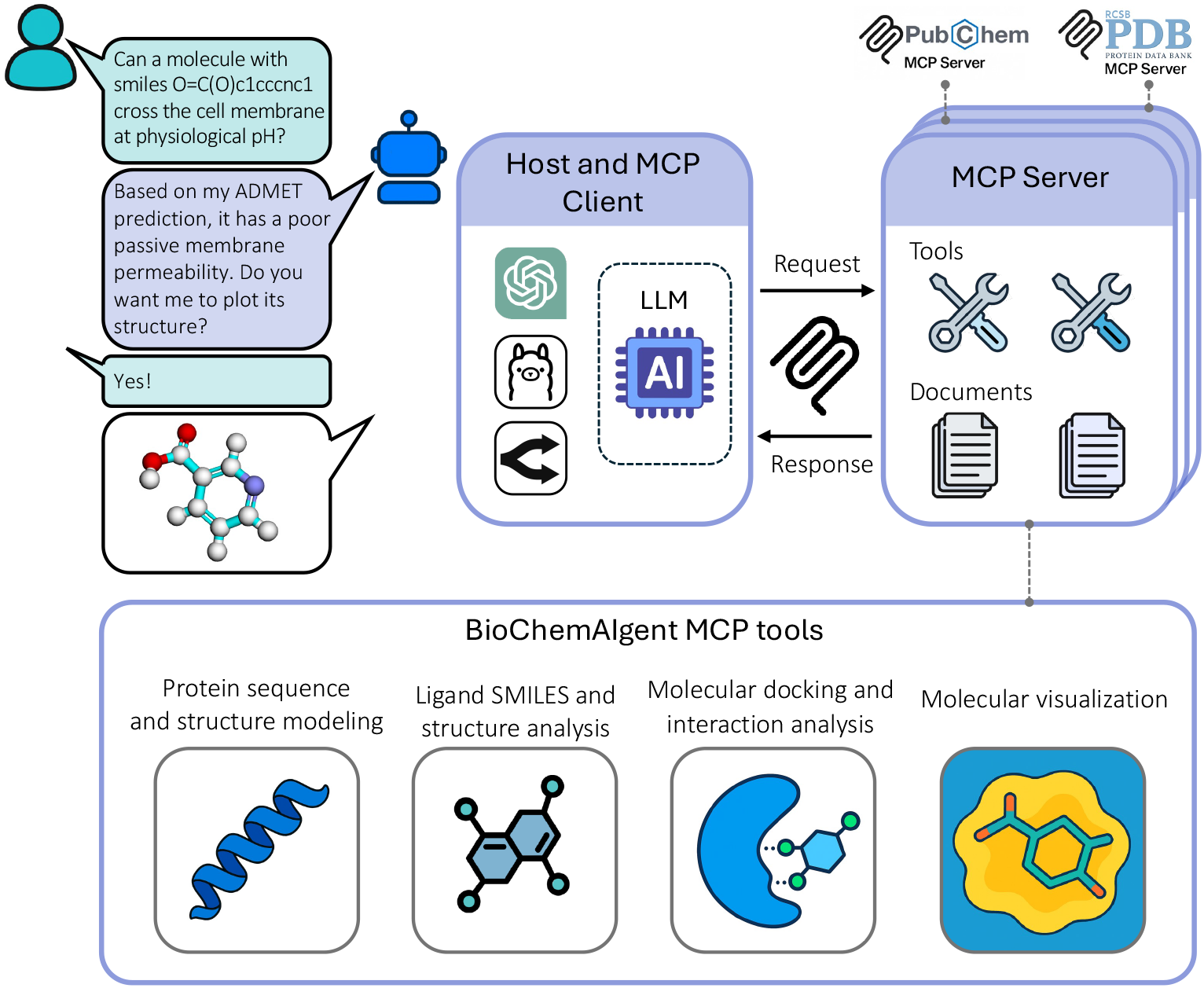
Overall architecture of BioChemAIgent. BioChemAIgent comprises a client, multiple servers, and a user-interface (UI) chatbot. The client follows the Model Context Protocol (MCP) and can be powered by large language models (LLMs) hosted by OpenAI, Ollama, or OpenRouter. Three MCP servers are integrated: PubChem-MCP-Server, PDB-MCP-Server, and a custom BioChemAIgent-MCP-Server equipped with tools for protein sequence and structure prediction and analysis, ligand SMILES and structure processing, molecular docking and interaction analysis, and molecular visualization. The server is also supplied with documentation that guides the agent toward best-practice use of these tools. An online UI is publicly accessible, providing both a conversational chatbot and a molecular structure viewer.

BioChemAIgent integrates 19 software tools and Python packages dedicated to biochemical analysis, structural biology, molecular docking and molecular visualization (Fig. 2; Supplementary Table 1). We note that certain tools are supported only when a dedicated user token is being provided.

**Figure 2.**
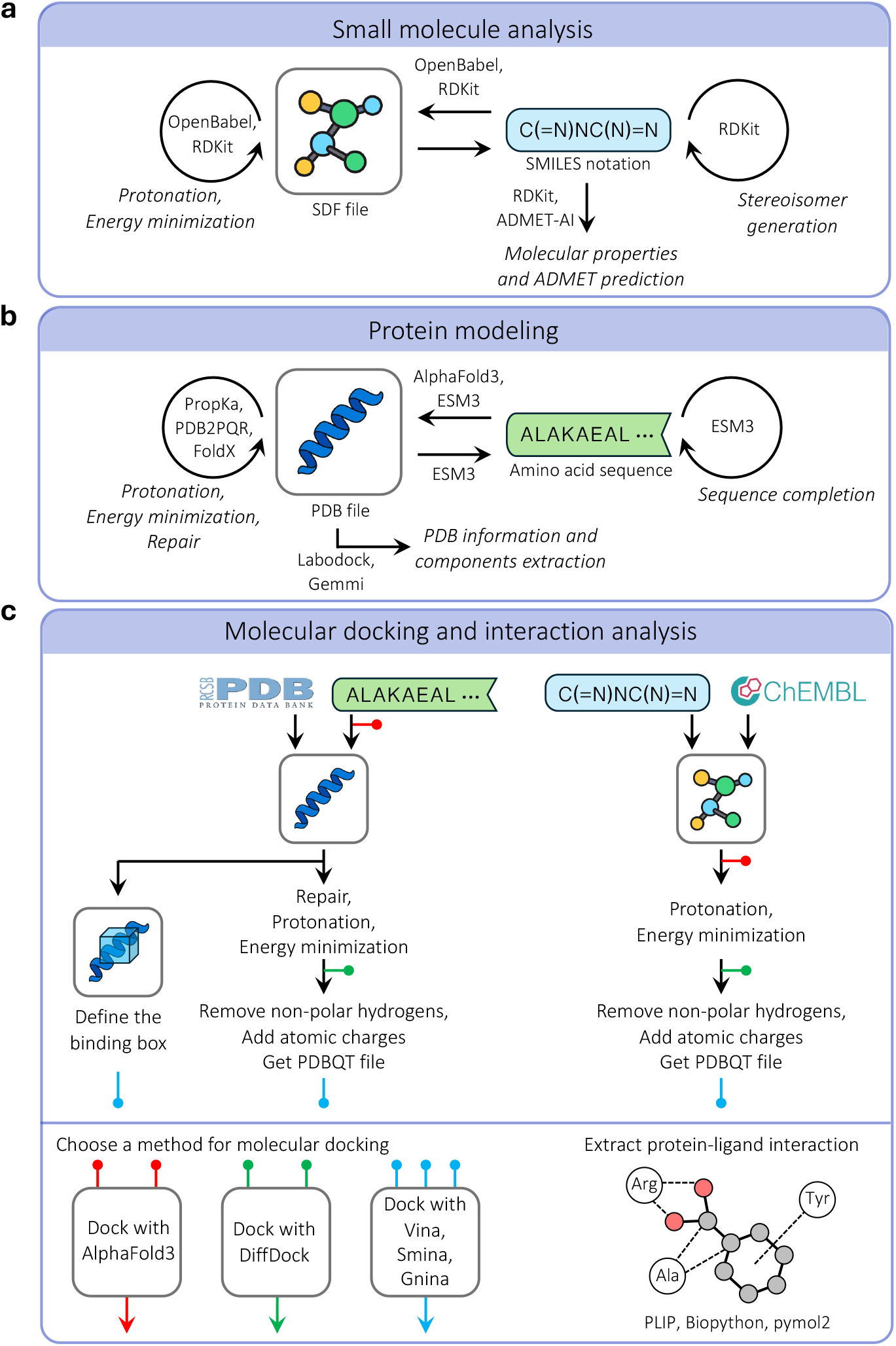
Schematic representation of BioChemAIgent workflows. (a) Small molecule analysis: analyzing ligand SMILES, 3D structure, given by structure data file (SDF), and converting them into each other. (b) Protein modeling: analyzing protein sequence, protein structure, given by protein data bank (PDB) file, and converting them into each other. (c) Molecular docking and interaction analysis: protein and ligand preparation, docking using different methods, i.e. Vina, Smina, Gnina, DiffDock, and AlphaFold 3, and protein-ligand interaction analysis.

- *Small molecule analysis*.Small molecules can be represented most efficiently by their SMILES notation which encodes their corresponding 3D structure. We developed MCP tools to convert these molecular representations into each other and to execute structure processing including energy minimization, protonation, and the generation of isomers, e.g., stereoisomers or tautomers. In addition, we incorporated tools to calculate molecular properties and predict ADMET parameters (i.e., absorption, distribution, metabolism, excretion, and toxicity; Fig. 2a).
- *Protein sequence and structure modeling*. Proteins are typically represented by their amino acid sequence or 3D structure as a PDB (protein data bank) file or an mmCIF (macromolecular crystallographic information file). Experimentally determined protein structures can be downloaded from RCSB PDB by the agent. For cases where experimental structures are not available, incomplete, or the resolution not sufficient, the agent is equipped with ESM3 ^2^ and AlphaFold3 ^3^, which are among the top AI-based protein structure prediction methods. BioChemAIgent uses ESM3 for protein sequence compilation, sequence prediction, 3D structure prediction, and function prediction. AlphaFold3 supports high-accuracy monomer and multimer structure prediction, and is also used for ligand docking in downstream workflows (see next). In addition, the agent can apply protein-processing steps, including protonation, repair, and energetic optimization, to prepare structures for downstream analysis (Fig. 2b).
- *Molecular docking and interaction analysis*. Molecular docking is a computational approach applied to evaluate the likelihood of ligand binding to a protein and to predict the ligand’s pose adopted upon binding, often accompanied by an estimate of the binding affinity. It is an essential step in drug discovery that guides hit identification, lead design, and optimization. BioChemAIgent integrates both physics-based docking methods, including AutoDock Vina ^9,10^, Smina, and Gnina ^11,12^, and deep learning-based methods, including DiffDock ^13^ and AlphaFold3, offering comprehensive assessment of ligand– receptor interactions. Physics-based methods require standardized preprocessing steps for both protein and ligand structures, including the removal of non-polar hydrogens, addition of partial atomic charges, and definition of the docking grid. Upon pose prediction, these methods then compute empirical scoring functions to approximate binding affinities and identify energetically favorable poses. For the analysis of protein-ligand complexes, we implemented a custom package to extract protein-ligand interactions, providing interpretable insights into molecular recognition mechanisms (Fig. 2c). BioChemAIgent selects from multiple docking and modeling backends (AutoDock Vina, Gnina, DiffDock, AlphaFold3) based on input type, use case, and resolution.
- *Molecular visualization*. To visualize molecular structures, complexes and their interactions, we developed two versatile and fully customizable tools that we introduce in this study: *render_structures* and *interaction_plot*. The *render_structures* function wraps *py3Dmol* (v 2.0.3) and provides 3D visualization of SDF (structure data file) and PDB files using user-defined style and surface rules. It also displays protein-ligand interactions, highlights docking grids, and supports applying specific styles to selected atoms or residues. The *interaction_plot* function provides a straightforward 3D view of protein-ligand interactions to facilitate presentation and interpretation of interactions.

To further provide accessibility to reliable online resources, we incorporated additional MCP servers for ChEMBL (PubChem-MCP-Server) and the Protein Data Bank (PDB-MCP-Server), enabling efficient retrieval of protein and compound information. For a more comprehensive list of MCP servers for biomedical research we refer the reader to the BioContextAI interface ^7^.

To showcase and evaluate the accuracy and fidelity of BioChemAIgent across drug discovery applications, we designed a comprehensive evaluation framework. This includes both LLM-based automatic and expert-based human assessment.

For *LLM-based automatic assessment*, a collection of expert curated questions and answers was prepared that probes different capabilities of BioChemAIgent (Supplementary Table 2). In particular, two types of ground-truth questions were considered including (i) tool call-based: those that require running a single tool and interpreting the result; and (ii) roadmap defining-based: those that require the agent to reason based on the available tools and provide a roadmap to achieve the goal. To automatically assess the agent’s ability in retrieving the correct answers, an LLM-based algorithm was developed (Supplementary Fig. 1). It first generates rephrased and corrupted versions of the original questions and labels agent’s answers as True, False, or Misinterpreted (i.e., problems in understanding the question). A comparison of the accuracy of BioChemAIgent responses powered by 10 different LLMs is shown in Fig. 3a and Supplementary Fig. 2.

**Figure 3.**
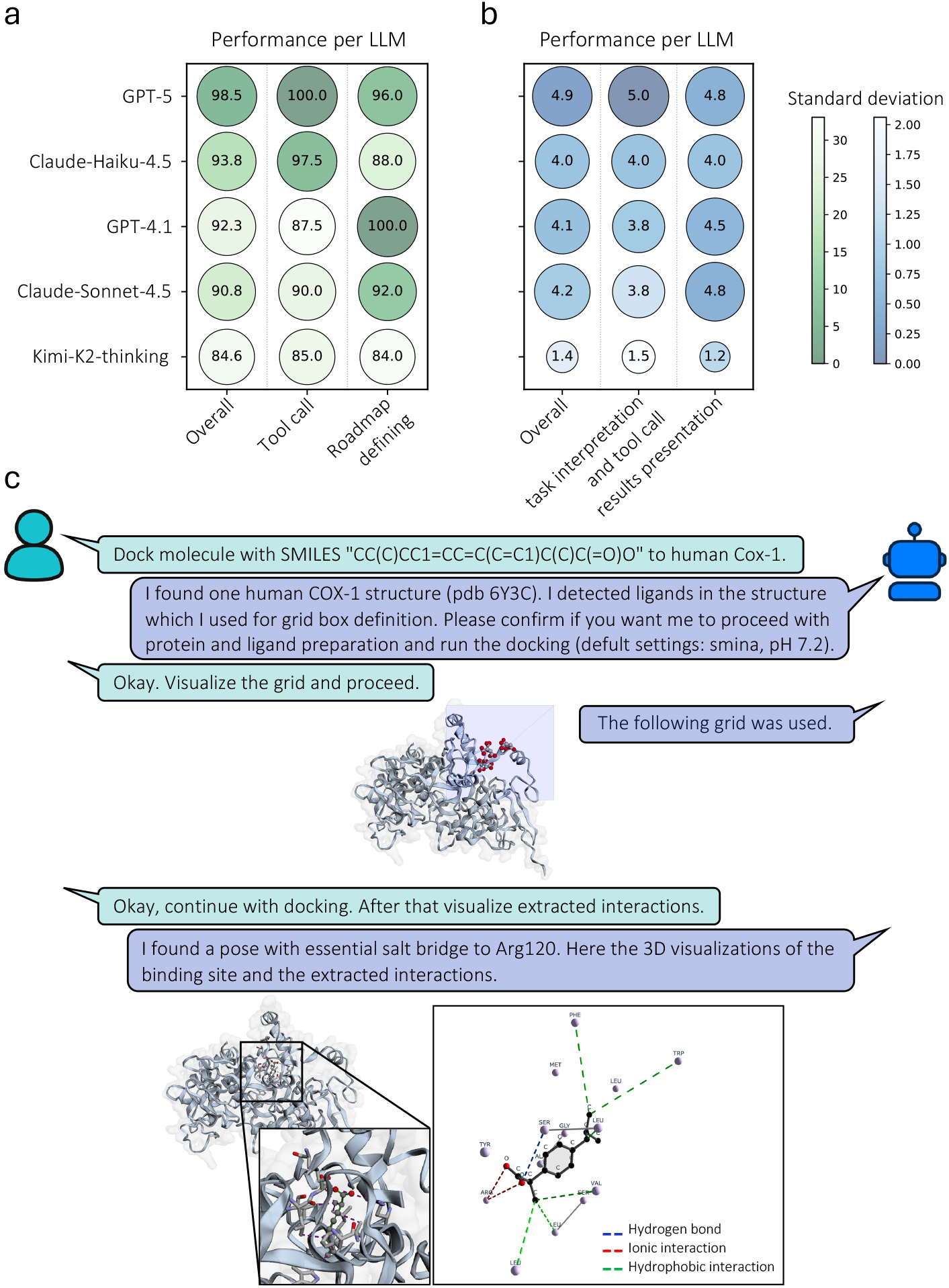
Evaluation and demonstration of BioChemAIgent. Performance comparison of (a) LLM-based automatic and (b) expert-based human assessment across different LLMs. Dot size and color intensity indicate the mean and standard deviation of scores across tasks. (c) A reduced illustration of a chat between user and agent for the task of molecular docking between Cox-1 and ibuprofen.

For *Expert-based human assessment*, the top 5 ranked LLMs from the automatic assessment were further evaluated through expert review. To this end, a domain expert designed four complex scenario-based tasks reflecting distinct realistic agent use cases (detailed in Supplementary Table 3-6). The expert evaluated the agent’s responses based on the correct use of tools and the accuracy in interpreting their output as well as on the precision in presenting the results and analysis. The agent received a score from 0 (failure) to 5 (perfect) for each criterion based on the expert’s prior defined expectations (Fig. 3b, Supplementary Table 7).

BioChemAIgent, particularly when powered by GPT-5, consistently met expert-level expectations across all evaluation criteria, including the formulation of actionable roadmaps for diverse drug-discovery tasks, correct tool use and accurate interpretation and presentation of output (Fig. 3a and 3b). In LLM-based automatic assessment, the agent successfully recovered ground-truth answers even when input queries were heavily corrupted in grammar and spelling. In expert-driven human evaluation, the agent was further challenged with a complex query, requiring implicit task inference rather than explicit instructions, yet it demonstrated the same high-level reasoning ability and strong domain knowledge in drug discovery (detailed in Supplementary Table 3-6). This result highlights that its performance did not merely rely on tool invocation, but on the agent’s ability to correctly interpret, contextualize, and reason over tool outputs in a manner comparable to domain experts.

Complete transcripts of the four original case study chats are available on the online platform ^i^. To showcase the performance of the GPT-5-powered BioChemAIgent, we curated a user-guided version of Case study 1 that explores molecular docking of ibuprofen to human Cox-1 (Fig. 3c). First, the agent queries UniProt for human COX-1 and on this basis successfully identifies the only matching PDB entry. After extracting its different components (i.e. protein chains, ligands, and solvent), it offers to define the docking grid’s localization and dimensions guided by co-crystallized ligand coordinates. By transparently considering case-specific parameters like the pH value and stereochemistry of the query ligand SMILES, the agent prepares both protein and ligand and executes docking with Smina by default. Leveraging domain knowledge, the agent recognizes an ionic interaction with Arg120 being essential for ibuprofen binding, and is even able to find it among the top-ranked poses. Finally, the agent visualizes the pose, and a 3D rendering of the extracted protein-ligand interactions completes the analysis by enabling rapid interpretation of the docking outcomes. Structure visualization is central to drug discovery, as it enables intuitive exploration and interpretation of drug mechanisms and interactions. While *py3Dmol* provides comprehensive rendering capabilities, meeting detailed specifications typically requires expert-level scripting. Our *render_structures* tool addresses this limitation by wrapping *py3Dmol* with an LLM-oriented manual, which allows users to describe their desired visualization in simple language. As a result, BioChemAIgent automatically renders and displays it in the user interface. Beyond Case study 1, we demonstrate a more complex visualization workflow in Case study 4 (Supplementary Fig. 3; Supplementary Table 6), that considers detailed specified visualization commands.

Overall, we believe that BioChemAIgent forms the foundation for community-oriented, agent-based *in silico* drug discovery. Its architecture supports dynamic communication between and across tools and curated documentation, and can naturally extend to future agent-to-agent infrastructures. We believe this design synergistically adds beyond the sum of its components, enabling workflow execution that matches expert-level analyses. BioChemAIgent was evaluated on expert-defined, complex drug-discovery tasks, where it fully satisfied the stated expectations. Our work further provides a community-oriented registry that enables more accessible, efficient, and reproducible drug discovery agents. We demonstrate how reasoning-capable agents can orchestrate domain-specific toolchains, paving the way for more accessible, scalable, and expert-level computational research in drug discovery and beyond.

## Online Methods

### Client design

To build the LLM client, we integrated OpenAI, OpenRouter and Ollama MCP services. They allow authenticated access to cloud-based state-of-the-art LLMs with tool support. While most options require usage-based charges, at the time of writing we identified Ollama’s *gpt-oss:120b-cloud* model as a free (but limited) option that performs well and offers access to BioChemAIgent at no cost. The client is connected to three servers: ChEMBL-MCP-Server, an MCP server providing advanced access to the ChEMBL chemical database ^14^, PDB-MCP-Server, an MCP server that provides access to the Protein Data Bank (PDB) ^15^, and BioChemAIgent-MCP-Server, developed in this study (see Server design section). When a user query is submitted, a combined message, including the system prompt, previous user-assistant exchanges, and the current query, together with the set of MCP tools are passed to the LLM. The response includes an assistant message and, optionally, a tool call. If a tool call is present, the client executes the specified tool and returns the result to the model, which then generates the final response. The client code is written in Python 3.11 using the OpenAI (v 1.75.0, with *AsyncOpenAI* class) and Ollama (v 0.6.0, with *Client* class) packages.

### Server design

BioChemAIgent-MCP-Server is composed of 27 tools for various drug-discovery tasks (Fig. 2), built on 19 software packages. Below, we describe the components and functionalities included in the current version of BioChemAIgent. A complete list of available tools is provided in Supplementary Table 1.

#### Small molecule analysis

Small molecules can be represented by their SMILES notation or their 3D structure representation (as SDF or PDB file). Certain small molecules can have stereocenters resulting in several stereoisomers. In this case, this information can either be provided by SMILES or it uses a tool based on RDKit ^16^ to generate all possible stereoisomers. To convert SMILES notation to a 3D structure, the BioChemAIgent performs energy minimization using OpenBabel (v 3.1.1). Another important step in generating the molecules’ 3D structure is protonation under certain environmental pH, which can further be done based on OpenBabel. Lastly, the agent can calculate molecules’ properties and and perform ADMET analysis, such as weight, LogP, TPSA, and number of atoms and bonds, using an RDKit-based tool. These properties are calculated based on RDKit and ADMET-AI ^17^ (Fig. 2a).

#### Protein sequence and structure analysis

ESM3 ^2^ and AlphaFold3 ^3^ represent the current flagships of AI-driven structural biology, trained on a vast corpora of protein sequences and structural data. ESM3 is a large-scale protein language model capable of bidirectional translation between amino acid sequences and three-dimensional structures, thereby capturing the fundamental principles of protein folding and function. AlphaFold3, in contrast, is a deep learning framework that extends structural prediction beyond individual proteins to accurately model complexes involving proteins, nucleic acids, ions, and small molecules, marking a significant advance in integrative structural biology. We implemented easy-to-use APIs for the use of these two models and integrated them with the MCP server. In addition, we implemented post-processing pipelines for protein protonation and structural optimization using propka ^18,19^, pdb2pqr ^20^, and FoldX ^21,22^, making them suitable for downstream analysis, such as molecule docking. FoldX identifies protein residues with poor torsion angles, van der Waals clashes, or unfavorable total energy, and then repairs them. It performs an optimization of side chains to remove minor van der Waals clashes and to locate new energy minima by exploring alternative rotamer configurations (Fig. 2b).

#### Molecular docking and interaction analysis

To estimate ligand-protein binding poses and binding affinities, we implemented a suite of state-of-the-art molecular docking tools. Molecular docking tools mainly can be categorized into physics-based approaches and machine learning-based methods. AutoDock Vina ^9,10^, Smina, and Gnina ^11,12^ represent physics-based methods, while DiffDock ^13^ and AlphaFold 3 represent deep learning-based methods (Fig. 2c). For ligands, these methods typically require an SDF file, which can be supplied directly or generated from a SMILES string. Ligands can be protonated at specific pH values and energy-minimized. For proteins, AlphaFold 3 only needs the amino-acid sequence, whereas other docking methods require the protein’s PDB file. Proteins can likewise be protonated at a chosen pH, repaired, and optimized. These steps prepare both molecules for DiffDock. AutoDock Vina, Smina, and Gnina require additional protein and ligand preparation, including removal of non-polar hydrogens and assignment of partial atomic charges. Users must also define a docking grid for docking, either covering the entire protein (blind docking) or restricted to known binding sites where native ligands are crystallized. These tools also compute scoring functions that estimate binding affinities. After docking and selecting a ligand pose, we used a custom package built on PLIP (v 2.3.0), Biopython (v 1.85), and pymol2 (v 3.1.0) to extract the protein-ligand complex, identify the binding pocket, and characterize protein-ligand interactions. The detected interactions include hydrogen bonds, hydrophobic contacts, halogen bonds, water bridges, salt bridges (ionic), π–π stacking, cation–π interactions, and metal coordination. We further developed dedicated visualization modules, using py3Dmol (v 2.0.3) and Plotly (v 6.3.0), to display these interactions.

### Agent instructions

To design an agent capable of performing complex tasks using a diverse set of provided tools, we identified two primary challenges. First, selecting the appropriate tool for a given task requires awareness of each tool’s advantages, limitations, and optimal application context. Second, implementing complete analytical pipelines, such as molecular docking, often demands the sequential integration of multiple of such specialized tools, each with distinct input formats and dependencies. To address these challenges, we implemented several key strategies.

#### System prompt design

The system prompt defines the agent’s behavior and scope. We augmented it with brief descriptions of available tools and workflows, together with explicit instructions for the agent to consult task-specific documentation (see below), generate an execution roadmap, and request user confirmation or modification before proceeding.

#### Structured documentation

For complex tools and workflows, we provided detailed instructional documents in markdown format. The agent is instructed to load and read each document prior to the tool’s first use, ensuring informed and context-aware execution.

#### Standardized output formatting

The output of each tool was converted into a structured format (python dictionary) that included: (i) the primary result; (ii) runtime messages such as success or warning indicators; and (iii) suggested next steps, where applicable. This standardized format enables communication between tools and facilitates automated pipeline execution.

### Performance evaluation

To evaluate the accuracy and fidelity of BioChemAIgent’s responses across multiple LLMs, two evaluation methods were designed and applied: LLM-based automatic assessment and expert-based human assessment. The LLM-based assessment enables large-scale evaluation across a broad set of questions and tests the robustness of BioChemAIgent to grammatical and spelling errors. In contrast, the expert-based human assessment provides an in-depth evaluation of BioChemAIgent’s performance on different drug discovery tasks by domain experts.

A schematic overview of the LLM-based automatic assessment is provided in Supplementary Fig. 1. First, a human expert curated 13 question-answer pairs covering diverse capabilities of BioChemAIgent (Supplementary Table 2). Next, an LLM (GPT-5.0) was employed to automatically generate 5 reformulated question variants, including both semantically rephrased and intentionally corrupted forms (system prompt in Supplementary Table 8). These LLM-generated queries were then provided to BioChemAIgent, instantiated separately with 10 different underlying LLMs, and the corresponding responses were collected. To evaluate the agent’s ability to retrieve the ground-truth answers, a separate LLM (GPT-5.0) was used to automatically assess whether each response contained the expected information (system prompt in Supplementary Table 8), labeling them as *True, False*, or *Misinterpreted* (i.e., problems in understanding the question). The evaluation results were further reviewed by an expert and refined where necessary. A response was considered to be *True* only when the agent’s output fully captured the ground-truth answer.

To enable an in-depth evaluation of BioChemAIgent, a human expert designed four case study tasks with predefined expected responses (Supplementary Tables 3–6). For each case study, BioChemAIgent was instantiated with one of five different LLMs, and its performance was independently evaluated by the expert. The assessment focused on A: the agent’s task interpretation, considering the correctness of tool call and accurate output interpretation, as well as on B: the agent’s transparency in explaining how the task was performed and its precision in results presentation. For each criterion A or B, the agent’s performance was rated with a score ranging from 0 (failure) to 5 (perfect), based on the expert’s predefined expectations.

## Code availability

The code for the user interface, MCP client, and MCP server, as well as agent instructions are available at https://github.com/imsb-uke/bcai. The GitHub repository also provides guidance for a community-oriented registry supporting the development of agentic systems for drug discovery and structural biology. In addition, The GitHub repository provides an easy-to-use Docker container that enables local installation and use of BioChemAIgent through the provided user interface. The user interface integrating the chatbot and molecular structure viewer is publicly accessible online at https://bcai.ims.bio; tools requiring academic access (listed in Supplementary Table 1) are available subject to possession of the appropriate licenses.

## Ethics declarations

Not applicable

## Competing interests

SWG, NCL, and SB work part time for Iniuva GmbH, a company developing AI-based drug discovery solutions. The authors declare no other conflicts of interest.

## Funding

This study was supported by grants from the Deutsche Forschungsgemeinschaft (DFG) to SB (SFB 1192 A2 and C3, TRR 422 CP2), LT (SFB 1700 SP01), and SH (SFB 1713 C01).

The project was funded by the Federal Ministry of Research, Technology and Space (Bundesministerium für Forschung, Technologie und Raumfahrt; BMFTR) as part of the German Center for Child and Adolescent Health (DZKJ) under the funding code 01GL2404A to SG.

## Authors’ Contributions

Conceptualization: BY and SB. Methodology & development: BY, NCL, SH, and LT. Data analysis: BY. Agent evaluation: BY and NCL. Writing original draft: BY and NCL. Reviewing and editing of the manuscript: BY, NCL, SG, SB. Visualization: BY and NCL. Supervision: SG and SB. Funding acquisition: SG and SB. All authors approved and contributed to the editing of the final draft.

## Acknowledgements

We would like to thank the members of the Institute of Medical Systems Bioinformatics for their feedback and Vadim Ustinov for IT support.

## Disclosure

During the preparation of this work the authors used ChatGPT to improve the creative writing process. Afterwards, the authors reviewed and edited the content as needed and took full responsibility for the content of the published article.

## Footnotes

i https://github.com/imsb-uke/bcai

ii https://bcai.ims.bio

i https://bcai.ims.bio

## Notes

### Summary of Updates

Here I updated the license from CC BY to CC BY-NC.

## References

1. Wei, H. & McCammon, J. A. Structure and dynamics in drug discovery. NPJ Drug Discov. 1, 1 (2024).

2. Hayes, T. et al. Simulating 500 million years of evolution with a language model. Science 387, 850–858 (2025).

3. Abramson, J. et al. Accurate structure prediction of biomolecular interactions with AlphaFold 3. Nature 630, 493–500 (2024).

4. Passaro, S. et al. Boltz-2: Towards accurate and efficient binding affinity prediction. bioRxivorg (2025) doi:10.1101/2025.06.14.659707.

5. Abdollahi, S. et al. A comprehensive comparison of deep learning-based compound-target interaction prediction models to unveil guiding design principles. J. Cheminform. 16, 118 (2024).

6. Hou, X., Zhao, Y., Wang, S. & Wang, H. Model Context Protocol (MCP): Landscape, security threats, and future research directions. arXiv [cs.CR] (2025).

7. Kuehl, M. et al. BioContextAI is a community hub for agentic biomedical systems. Nat. Biotechnol. 43, 1755–1757 (2025).

8. Tang, L. Artificial intelligence agents for biology. Nat. Methods 22, 2496–2497 (2025).

9. Eberhardt, J., Santos-Martins, D., Tillack, A. F. & Forli, S. AutoDock Vina 1.2.0: New docking methods, expanded force field, and Python bindings. J. Chem. Inf. Model. 61, 3891–3898 (2021).

10. Trott, O. & Olson, A. J. AutoDock Vina: improving the speed and accuracy of docking with a new scoring function, efficient optimization, and multithreading. J. Comput. Chem. 31, 455– 461 (2010).

11. McNutt, A. T. et al. GNINA 1.0: molecular docking with deep learning. J. Cheminform. 13, 43 (2021).

12. McNutt, A. T., Li, Y., Meli, R., Aggarwal, R. & Koes, D. R. GNINA 1.3: the next increment in molecular docking with deep learning. J. Cheminform. 17, 28 (2025).

13. Corso, G., Stärk, H., Jing, B., Barzilay, R. & Jaakkola, T. DiffDock: Diffusion steps, twists, and turns for molecular docking. arXiv [q-bio.BM] (2022).

14. ChEMBL-MCP-Server: A Comprehensive Model Context Protocol (MCP) Server Providing Advanced Access to the ChEMBL Chemical Database. (Github).

15. PDB-MCP-Server: A Model Context Protocol (MCP) Server That Provides Access to the Protein Data Bank (PDB) - the Worldwide Repository of Information about the 3D Structures of Proteins, Nucleic Acids, and Complex Assemblies. (Github).

16. Landrum, G. RDKit. https://www.rdkit.org.

17. Walther, P. et al. ADMET-AI: A Machine Learning ADMET Platform for Evaluation of Large-Scale Chemical Libraries Open Access Kyle Swanson.

18. Søndergaard, C. R., Olsson, M. H. M., Rostkowski, M. & Jensen, J. H. Improved treatment of ligands and coupling effects in empirical calculation and rationalization of pKa values. J. Chem. Theory Comput. 7, 2284–2295 (2011).

19. Olsson, M. H. M., Søndergaard, C. R., Rostkowski, M. & Jensen, J. H. PROPKA3: Consistent treatment of internal and surface residues in empirical pKa predictions. J. Chem. Theory Comput. 7, 525–537 (2011).

20. Jurrus, E. et al. Improvements to the APBS biomolecular solvation software suite. Protein Sci. 27, 112–128 (2018).

21. Delgado, J. et al. FoldX force field revisited, an improved version. Bioinformatics 41, (2025).

22. Delgado, J., Radusky, L. G., Cianferoni, D. & Serrano, L. FoldX 5.0: working with RNA, small molecules and a new graphical interface. Bioinformatics 35, 4168–4169 (2019).

