## supplementary materials for "BioChemAIgent: An AI-driven Protein Modeling and Docking Framework for Structure-Based Drug Discovery" (from another version of this preprint: version 1, 2025-12-19)

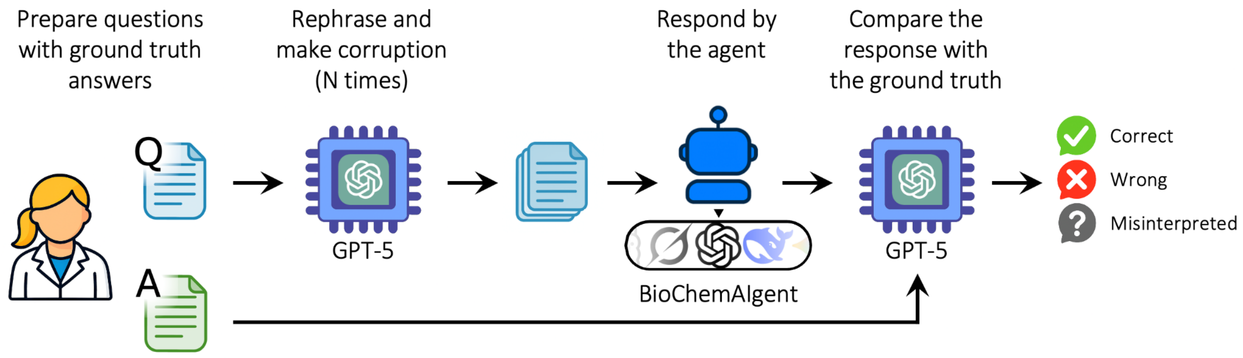


**Supplementary Figure 1. LLM-based automatic assessment.** A set of questions and ground-truth answers are prepared by an expert. The questions are rephrased and corrupted by an LLM (GPT-5.0) generating 5 rephrased variants per question. The questions are then provided to BioChemAIgent, powered by different LLMs, and the responses were fed to a second LLM (GPT-5.0) to be compared with the ground-truth answers labeling them as *True*, *False*, or *Misinterpreted*.

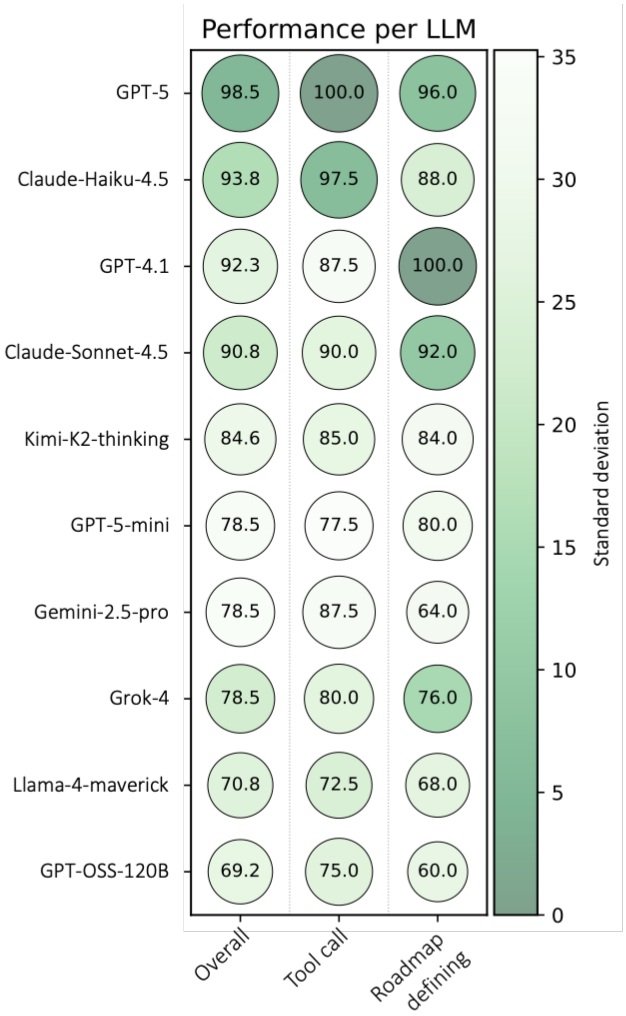


**Supplementary Figure 2. LLM-based automatic assessment across 10 different LLMs.** Two groups of ground-truth questions were considered including those based on (i) tool call and results interpretation, and (ii) roadmap definition.

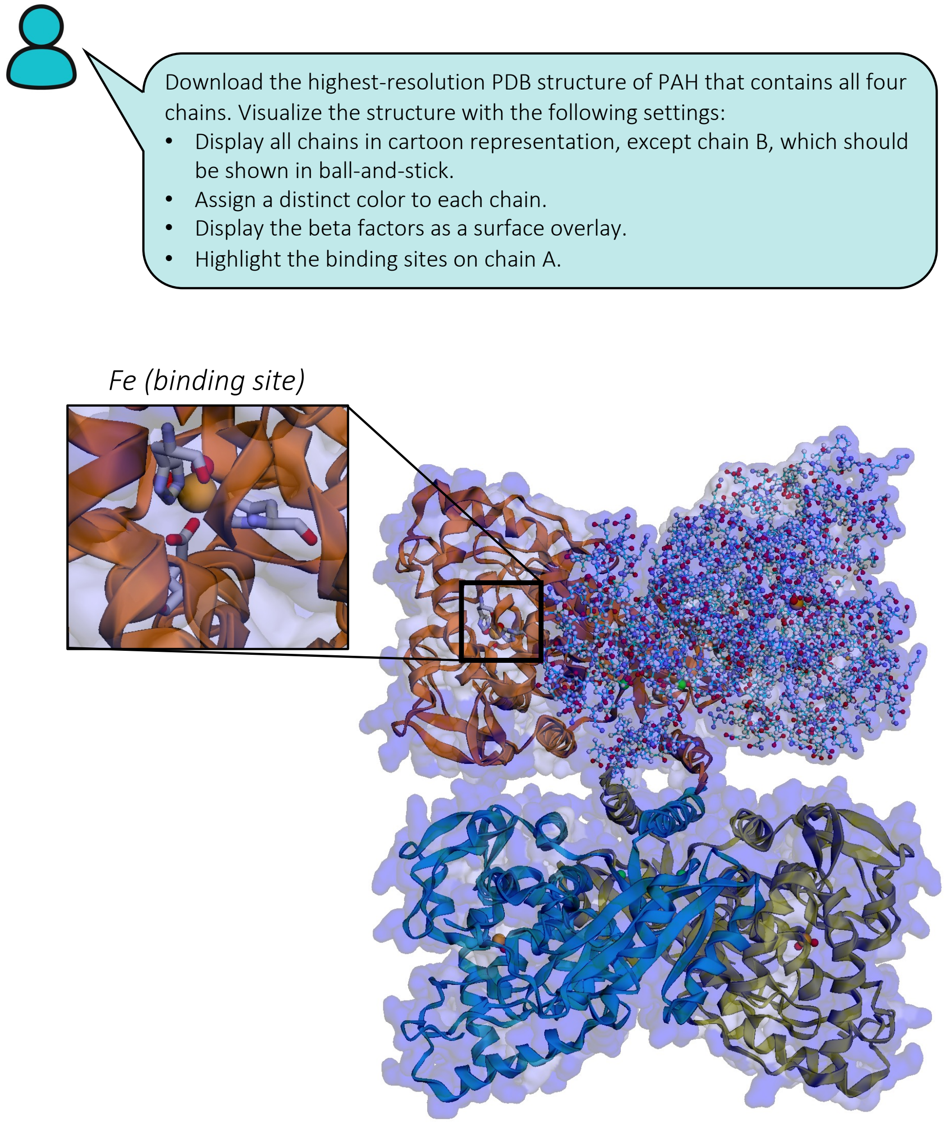


**Supplementary Figure 3. Visualizing result for Case study 4.** Downloading a specified PDB file and visualizing it according to user-defined requirements, performed by the GPT-5-powered agent.

**Supplementary Table 1.** Software and Python packages integrated by BioChemAIgent

| Analysis type | Software | Version | Use Case | License |
|---|---|---|---|---|
| Ligand and SDF file processing | RDKit | 2022.9.5 | Ligand-based analysis and properties prediction | Open-source |
| | ADMET-AI | 1.4.0 | ADMET prediction | Open-source |
| | OpenBabel | 3.1.1 | Ligand protonation and energy minimization | Open-source |
| Protein and PDB file processing | labodock | 2.0.0 | PDB file processing | Open-source |
| | gemmi | 0.7.3 | PDB and mmCIFfile data extraction | Open-source |
| | propka | 3.5.1 | Protein protonation | Open-source |
| | pdb2pqr | 3.6.1 | Protein protonation and structural optimization | Open-source |
| | FoldX | 5.1 | Protein repair and structural optimization | Academic use |
| | ESM3 | 3.2.1 | Protein sequence and structure prediction | Academic use (via user token) |
| | AlphaFold3 | 3.0.1 | Protein and ligand structure prediction | Academic use (via user token) |
| Molecular Docking | Autodock Vina | 1.2.5 | Physics-based molecular docking | Open-source |
| | Smina | 2024.7.10 | Physics-based molecular docking | Open-source |
| | Gnina | 1.0 | Physics-based molecular docking | Open-source |
| | DiffDock | 1.1 | Deep learning-based molecular docking | Open-source |
| Protein-ligand interaction | PLIP | 2.3.0 | Protein-ligand interaction analysis | Open-source |
| | Biopython | 1.85 | Protein-ligand interaction analysis | Open-source |
| | pymol2 | 3.1.0 | Protein-ligand complex file processing | Open-source |
| Visualization | py3Dmol | 2.0.3 | Protein and ligand visualization | Open-source |
| | Plotly | 6.3.0 | Interaction visualization | Open-source |

**Supplementary Table 2.** The list of expert-curated questions and answers for LLM-based automatic assessment. We consider two types of questions: Tool-based questions and reasoning-based. For the tool-based questions, we add the following to the end of the questions: "*Do NOT provide any roadmap. Do NOT ask for confirmation of the roadmap. Directly run the tools and write down the result.*"

| Q/A No. | Type | Content |
|---|---|---|
| Q1 | Tool call | What is the smiles of BH4 |
| A1 | | C[C@H](O)[C@H](O)[C@H]1CNc2nc(N)[nH]c(=O)c2N1, or<br>CC(O)C(O)C1CNc2nc(N)[nH]c(=O)c2N1 |
| Q2 | Tool call | Generate all stereoisomers of CC(O)C(O)C1CNc2nc(N)[nH]c(=O)c2N1 |
| A2 | | Iso1: C[C@H](O)[C@H](O)[C@H]1CNc2nc(N)[nH]c(=O)c2N1<br>Iso2: C[C@H](O)[C@H](O)[C@@H]1CNc2nc(N)[nH]c(=O)c2N1<br>Iso3: C[C@H](O)[C@@H](O)[C@H]1CNc2nc(N)[nH]c(=O)c2N1<br>Iso4: C[C@H](O)[C@@H](O)[C@@H]1CNc2nc(N)[nH]c(=O)c2N1<br>Iso5: C[C@@H](O)[C@H](O)[C@H]1CNc2nc(N)[nH]c(=O)c2N1<br>Iso6: C[C@@H](O)[C@H](O)[C@@H]1CNc2nc(N)[nH]c(=O)c2N1<br>Iso7: C[C@@H](O)[C@@H](O)[C@H]1CNc2nc(N)[nH]c(=O)c2N1<br>Iso8: C[C@@H](O)[C@@H](O)[C@@H]1CNc2nc(N)[nH]c(=O)c2N1 |
| Q3 | Tool call | Generate the 3D structure of CC(=O)Oc1ccccc1C(=O)O with MMFF94 force field. |
| A3 | | Tool call, name:smiles_to_3d, inputs: ligand_smiles = “CC(=O)Oc1ccccc1C(=O)O”, force_field = “MMFF94” |
| Q4 | Tool call | Protonate the ligand SDF file at files/queried_ligand.sdf at pH=7.1 |
| A4 | | Tool call, name:protonate_and_optimize_ligand, inputs: input_file = files/queried_ligand.sdf, ph = 7.1 |
| Q5 | Tool call and reason | Will the caffeine with SMILES: "CN1C=NC2=C1C(=O)N(C)C(=O)N2C" and aspirin with smiles “CC(=O)Oc1ccccc1C(=O)O” pass the cell membrane? |
| A5 | | Caffeine yes; aspirin limited at pH=7 but better at lower pH |
| Q6 | Tool call and reason | What is the hydrophobicity of the ligand with the smiles of "C[C@H](O)[C@H](O)[C@H]1CNc2nc(N)[nH]c(=O)c2N1" |
| A6 | | clogP = −1.70, strongly hydrophilic, not hydrophobic |
| Q7 | Roadmap defining | Visualize the 3D structure of protein sequence: ALEKALAEKALA. |
| A7 | | Here is the roadmap:<br>1. Predict the 3D structure with ESM3 or AlphaFold3;<br>2. Make the visualization of the generated 3D structure. |

**Supplementary Table 2 (cont.)**

| Q/A No. | Type | Content |
|---|---|---|
| Q8 | Roadmap defining | Prepare this ligand Cl.OC[C@H]1NC[C@H](O)[C@@H](O)[C@H]1O for docking. |
| A8 | | Here is the roadmap:<br>1. Create sdf of this exact isomer removing the counter ion.<br>2. Protonate the ligand at the desired pH (optional).<br>3. For Vina/Smina/Gnina: create PDBQT file<br>4. Make docking query file |
| Q9 | Roadmap defining | Prepare Alpha-Galactosidase for docking. |
| A9 | | Here is the roadmap:<br>1. Use ChEMBL to find pdb id<br>2. Download the pdb file<br>3. Extract components<br>4. Protonation and optimization/energy minimization at the desired pH<br>5. Choose the docking site (full protein or binding site of the co-crystallized ligand) |
| Q10 | Tool call | Predict the most probable missing amino acids in the sequence ALEK_L_EK_LA. |
| A10 | | Tool call, name:run_esm3, inputs: protein_input = {'sequence' : ALEK_L_EK_LA}, task = “Sequence” |
| Q11 | Tool call and reason | What is the resolution of the pdb file at “files/queried_protein.pdb”. Is that suitable for docking? |
| A11 | | Resolution: 3.06 Å (X-ray diffraction), it is moderate to low. |
| Q12 | Roadmap defining | What are the charge states of a ligand, given its SMILES, to have at physiological or acidic conditions, respectively? |
| A12 | | Here is the roadmap:<br>1. Convert to sdf by running protonation at pH 7.0 or 2.0<br>2. Calculate the net charge of the molecule. |
| Q13 | Roadmap defining | Visualize Hemoglobin |
| A13 | | Here is the roadmap:<br>1. Search for the pdb ids of Hemoglobin.<br>2. Download the desired pdb file<br>3. Make the HTML visualization |

**Supplementary Table 3.** Human expert evaluation for Case study 1.

| Tite | Content |
|---|---|
| Case study 1 | Dock CC(C)CC1=CC=C(C=C1)C(C)C(=O)O to human Cox-1 |
| Expert's expectation | This task assesses the agent's performance on a molecular docking-related task in a realistic drug discovery setting. (i) The ligand has a stereocenter; the agent should notice it and ask with which enantiomer to proceed. It would also be a bonus if the agent by itself recognizes that the ligand is ibuprofen. (ii) The agent should download the only available human Cox-1 crystal structure (PDB ID: 6Y3C) and extract its components. (iii) The crystal structure contains non-functional ligands, which should be noticed by the agent. Accordingly, the agent has to suggest that the docking grid should contain the whole protein instead of defining it around the non-functional ligands. (iv) It should ask for the docking method, offer to change the pH, prepare protein and ligand for docking (protonation, energy minimization), run the docking, and list affinity scores of the top poses. (v) Finally it should offer to extract the protein-ligand interactions and suggest visualization of the complex. The agent receives full credit only if it handles all of these elements correctly with proper explanation. |
| Expert's evaluation | The agent based on GPT-5 correctly executed all the steps and was the only model to recognize that the SMILES code corresponds to ibuprofen. Even though only one human PDB structure was available, the agent noted its general preference for holo structures in docking, which indeed would be optimal. By explicitly and transparently outlining the steps it performed for the preparation of both the ligand and the protein, the agent provided valuable guidance for the user to investigate the results. Outstanding was the agent's consideration of the essential ionic interaction with Arg120. Agents based on other LLMs were able to successfully finish the task by using the correct PDB; however, not all of them identified and used the UniProt ID and therefore considered wrong proteins as well, without giving a clear warning that they are not human or even distinct proteins (GPT-4.1 and Claude-Haiku-4.5). Moreover, correct recognition of other molecules present in the PDB structure as being non-functional components was unique to the agent based on GPT-5 and therefore the other agents did not consider this for grid definition. Some agents even did not transparently disclose the grid's localization (Claude-Sonnet-4.5, Claude-Haiku-4.5). A clearly visible warning for chirality of the ligand was exclusive to agents based on GPT-5 and Claude-Sonnet-4.5, whereas the others randomly selected one of the two enantiomers without specification or further notice. The presentation of interim results and used input features were less precise for agents based on Claude-Sonnet-4.5 and GPT-4 compared to the others, while the Claude-Sonnet-4.5-based agent noticeably required a higher number of confirmations by the user, which made its handling less user-friendly. Because the kimi-k2-thinking–based agent did not produce a usable response within a reasonable timeframe, its response was excluded from evaluation and it was assigned 0/5 for both criteria A and B. |

**Supplementary Table 4.** Human expert evaluation for Case study 2.

| **Task** | **Content** |
|---|---|
| Case study 2 | Will the caffeine with SMILES: "CN1C=NC2=C1C(=O)N(C)C(=O)N2C" and aspirin with smiles "CC(=O)Oc1ccccc1C(=O)O" pass the cell membrane? |
| Expert's expectation | This task evaluates the agent's domain knowledge of biochemistry and molecular properties related with drug-likeness. The short answer should be: caffeine will pass the cell membrane at physiological pH; aspirin exhibits poor membrane permeability at neutral pH, which will be markedly increased in the acidic environment of the stomach. To reach this conclusion, the agent should calculate logP, topological polar surface area (TPSA), molecular weight, number of rotatable bonds, hydrogen bond donors (HBD), hydrogen bond acceptors (HBA) and consider pKa values. Both compounds fulfill Lipinski rules. However, in particular, aspirin will be charged at neutral pH, which is associated with poor passive permeation. |
| Expert's evaluation | All the agents except for the one based on Kimi-K2-thinking, were able to call the correct tool required for this task, which is either admet_predict or calculate_ligand_properties calculation. The GPT-5-powered agent even used both tools and considered their output for solving the task. However, only the GPT-5-powered agent considered the pKa values and thus correctly responded to the question. Moreover, it consolidated publicly available data on aspirin's and caffeine's membrane permeability for the final evaluation which supported its calculated values and interpretation and therefore resulted content-wise in an outperforming response to this question. However, the agent presented the results and explanation less well-structured than agents based on Claude-Sonnet-4.5 and GPT-4 did. The agent based on Claude-Haiku-4.5 presented an incomplete set of calculated values and omitted properties essential to answering the question. |

**Supplementary Table 5.** Human expert evaluation for Case study 3.

| Task | Content |
|---|---|
| Case study 5 | Visualize the protein with sequence AEKLAEALA??A |
| Expert's expectation | This task evaluates the agent's ability in implicit task inference, while assessing its performance on a complex protein modeling application. The agent should infer the required steps: (i) complete the sequence first, (ii) then model the 3D structure, [iii] finally visualize it. The only tool available for the first task is ESM3, while for the second task, both ESM3 and AlphaFold 3 can be used. Therefore, the agent is expected to run ESM3 for the first step and then ask to choose between ESM3 and AlphaFold3 for the second task. Lastly, it should visualize the predicted structure. The agent receives full credit only if it handles all of these elements correctly with proper explanation. |
| Expert's evaluation | The agents powered by all LLM models were able to successfully complete the task as expected. However, the Claude-Sonnet-4.5-based agent required a noticeably higher number of user confirmations, similar to what was observed in Case study 1. The same was observed for kimi-k2-thinking, where additionally cryptic phrasing in its responses impeded smooth user interaction. |

**Supplementary Table 6.** Human expert evaluation for Case study 4.

| Task | Content |
|---|---|
| Case study 5 | Download the highest-resolution PDB structure of PAH that contains all four chains. Visualize the structure with the following settings:<br>● Display all chains in cartoon representation, except chain B, which should be shown in ball-and-stick.<br>● Assign a distinct color to each chain.<br>● Display the beta factors as a surface overlay.<br>● Highlight the binding sites on chain A. |
| Expert's expectation | This task involves multiple interdependent steps, requiring the agent to select appropriate tools and craft precise prompts for each stage, a process that would be substantially time-consuming for a human to perform manually. The agent should first retrieve all PDB entries for PAH, extract their metadata, and identify the structure that contains four chains with the highest available resolution. The *render_structures* tool should then be used to produce the visualization, which requires domain knowledge in *py3Dmol* to meet all rendering specifications. Finally, the agent should highlight the binding sites in chain A, using either its domain knowledge or information obtained by executing available tools. |
| Expert's evaluation | Only the GPT-5-based agent completed the task flawlessly, without requiring any user intervention (Supplementary Fig. 3). It used online databases to retrieve the protein information to find the best matching PDB ID, plotted the structure as required, and inferred the binding sites from its internal knowledge. The GPT-4-based agent initially encountered an error when retrieving protein information but solved it on its own and ultimately selected the correct PDB ID. It produced the structure plot, though it required an additional iteration to refine it. For the binding site, it added only a label at the iron-binding position. Both the Claude-Sonnet-4.5-based and Claude-Haiku-4.5-based agents misinterpreted the protein name and needed the user to supply the complete name. To identify the PDB ID, the former downloaded a random selection of PDB files to inspect them, an inefficient strategy, while the latter queried an online database directly. Each produced the required structural plot and marked the iron-binding site with a label. After a follow-up request on highlighting the binding sites, both used a tool to extract the residues at the binding site and added them correctly to the plot. The Kimi-K2-thinking-based agent retrieved a PDB ID without explaining its reasoning, and the final structure plot only partially met the expected criteria. |

**Supplementary Table 7.** Expert rating of BioChemAIgent powered by different LLMs. Possible scores range from 0 (failure) to 5 (perfect) for each criterion (A: task interpretation and tool call, B: transparency of explanation and precision in results presentation) and were given based on the expert's prior expectations.

| Case study | Criterion | GPT-5 | GPT-4.1 | claude-sonnet-4.5 | claude-haiku-4.5 | kimi-k2-thinking |
|---|---|---|---|---|---|---|
| 1 | A | 5 | 3 | 5 | 4 | 0 |
| | B | 5 | 4 | 4 | 4 | 0 |
| 2 | A | 5 | 4 | 3 | 3 | 0 |
| | B | 4 | 5 | 5 | 3 | 1 |
| 3 | A | 5 | 5 | 5 | 5 | 5 |
| | B | 5 | 5 | 5 | 4 | 3 |
| 4 | A | 5 | 3 | 2 | 4 | 1 |
| | B | 5 | 4 | 5 | 5 | 1 |

**Supplementary Table 8.** System prompts used for the LLMs in the automatic assessment workflow.

| Task | System prompt |
|---|---|
| Rephrase and make corruption | Rephrase and intentionally corrupt the given text by adding typos and grammatical errors so that the meaning becomes somewhat unclear.<br>Add no extra text. Only write down the rephrased text.<br>You can use words with similar meaning, for example: ligand = small molecule, protein = polypeptide, ...<br>Do not corrupt the SMILES notation if any.<br><br>**Examples**<br>User: The ligand is hydrophobic.<br>Agent: the smal mulecul avoids whater. |
| Compare the response with the ground truth | You need to compare the two given texts.<br><br>- A is the ground truth text written by an expert.<br>- B is a text generated by AI.<br>- Check if the text B has the information aligned with A.<br>- If yes, answer "True", if not, answer "False".<br>- If the text B indicates that the LLM did not understand the question, then answer "Misinterpreted"<br>- Your answer should only be one word among: True, False, Misinterpreted<br><br>**Examples**<br>A: The ligand is hydrophobic.<br>B: The small molecule avoids water.<br>Answer: True<br><br>A: The ligand binds the protein.<br>B: The docking is not strong.<br>Answer: False<br><br>A: The ligand binds the protein.<br>B: Please repeat the question.<br>Answer: Misinterpreted |
